# A multi-tissue atlas of age-associated differential transcript usage in mouse reveals predominantly regulatory isoform switching across brain, liver, and adipose

**DOI:** 10.64898/2026.11.10.765779

**Authors:** Nishitha Paidimukkala, Syed S Ali, Linshan Laux, Ann V. Hertzel, Mythili Dileepan, Erich Kumerfeld, Constantin Aliferis, Paul D. Robbins, Laura J. Niedernhofer, Jinhua Wang

## Abstract

Alternative splicing is increasingly recognized as dysregulated in the aging process across tissues and affects proteome and other regulatory changes which can be biomarkers of aging. In this study, we aimed to identify differential transcript usage (DTU) and isoforms switch accompanied with functional annotation and structural analysis in mouse brain, liver and adipose tissue during natural aging and Ercc1−/Δ mice. Age associated isoform switches affected both coding and non-coding transcript features. Across tissues, 36.5–42.6% of switches altered the predicted coding sequence, 31.2–36.5% involved non-coding alternative arms, and 11–18% involved transcripts encoding an identical protein. We observed overall 3’UTR lengthening in the brain and shortening in the adipose. A single pan-organ isoform switch event in the gene *Ighm*, recurs in seven separate tissue samples. BioEMU modeling confirms feasibility for a minority of switches following a seven-fold eligibility cascade; the *Cyth1* exemplar shows an intrinsically disordered N-terminal extension that does not change the shared SEC7 and PH domains. Together, these results distinguish age-associated isoform changes as predominantly regulatory in mechanism, with structural consequences restricted to tissue-specific events that form a tractable target for conformational ensemble modeling.

## Introduction

The relative abundance of transcript isoforms can change with age, with potential consequences for protein sequence and RNA regulation. These changes can occur even when overall gene expression remains unchanged. Differential transcript usage (DTU) analysis quantifies these changes in the relative proportions of a gene’s transcript isoforms, identifying age associated shifts that may occur without changes in overall gene expression. However, it remains unclear which transcripts shift during aging and whether these switches are shared across organs tissues and how often they alter the encoded protein or non-coding features of the transcript.

Previous studies have identified age-associated splicing changes across tissues and shown that specific isoforms can regulate cellular senescence. Rodríguez et al.^1^ identified changes in exon usage in mouse tissues and reported enrichment of RNA-processing functions and the spliceosome pathway among the affected genes. Sieber, Barth, and Marz^2^, however, described a largely stable splicing landscape across tissues and species, with approximately 80% of genes undergoing a dominant-isoform switch retaining the same number of encoded protein domains. Studies of individual genes show that isoform changes can directly influence cellular senescence. The long LEF1 isoform protects against cellular senescence.^3^ Opposing effects have also been demonstrated for p53 isoforms: Δ133p53 suppresses replicative senescence, whereas p53β promotes it.^4^ The mitochondrial GRSF1 isoform also declines in senescent cells in aged mouse skeletal muscle and heart, but not in liver, indicating tissue-dependent changes.^5^

Studies of individual genes have established the functional importance of specific isoforms, while transcriptome-wide analysis can reveal the extent and consequences of isoform switching across aging tissues. Moreover, exon-level measurements and conserved protein-domain counts do not resolve whether individual switches alter coding sequences, untranslated regions, or both. Comparing transcript pairs across organs is therefore needed to distinguish shared from tissue-specific switches and assess their predicted functional consequences.

We did a pan tissue study to identify transcript isoform shifts across individual genes and functionally annotated the significant isoforms to understand the changes across multiple organ systems. Here, we examined how transcript isoform proportions change with age across mouse brain, liver, and adipose tissue, including natural-aging cohorts and liver from Ercc1−/Δ mice. We applied DRIMSeq to test for differential transcript usage and stageR to control the false discovery rate at the gene and transcript levels. Within individual genes, we identified transcript pairs in which one isoform increased in relative usage while the other decreased, defining these events as significant isoform switches. We quantified the magnitude of these changes and examined whether the same transcript pairs switched across tissues. To assess their potential functional consequences, we compared annotation matched transcripts for differences in coding potential, predicted protein domain sequence, and length of 5′ and 3′ untranslated regions. We then tested whether switch genes were enriched for biological processes or pathways, using genes tested for differential transcript usage as the background.

Our analyses showed that age-associated isoform switching affects both coding and noncoding transcript features, with distinct patterns across the tissues we included in this study. Significant isoforms gained relative abundance within aging tissue showed longer 3′UTRs in brain and shorter 3′UTRs in adipose tissue, suggesting tissue-dependent changes in transcript regulation. Selected switches also recurred in Ercc1−/Δ mice, indicating partial overlap with natural aging.

Structural ensemble modeling of protein-altering switches further allowed us to examine whether sequence differences affected the shared protein fold. Together, these analyses connect changes in transcript proportions to their predicted functional consequences across aging tissues.

## Results

### Cross tissue survey

To characterize age-associated changes in transcript composition, we analyzed RNA-seq data from mouse brain, liver and adipose tissue across 13 young-versus-aged comparisons (Fig. 1; Table 1). We examined seven brain regions from female mice aged 3 and 24 months, together with a separate midbrain cohort containing both sexes. In the liver, we compared male mice aged 16 weeks, 1.5 and 2 years with an 8-week-old reference group. The adipose analysis included three depots from male mice: epididymal, inguinal and mesenteric.

**Figure 1.**
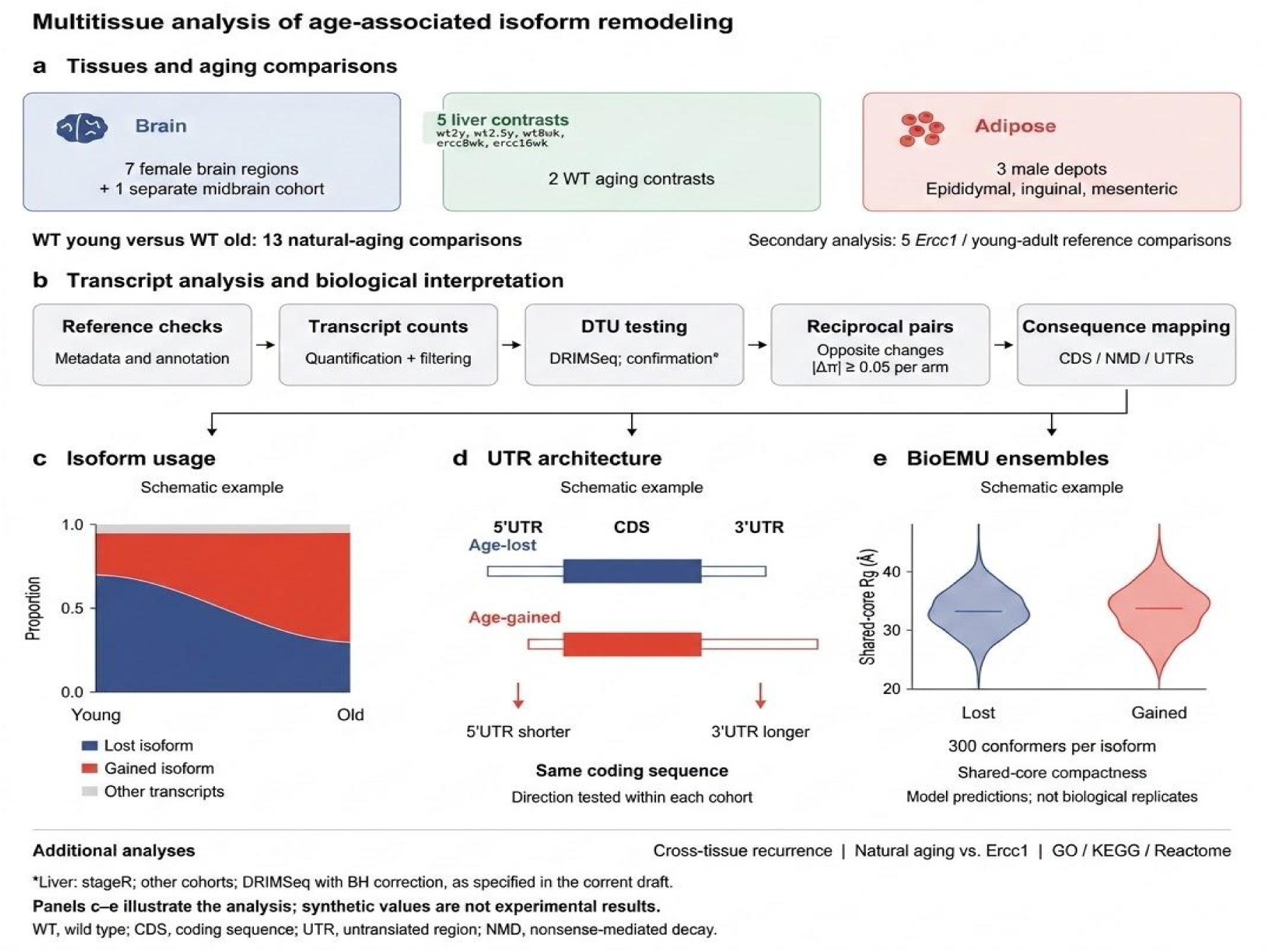
Study design and analysis pipeline. Overview of the study design across brain, liver, and adipose tissues, depicting the natural aging and *Ercc1*−/Δ cohorts alongside the transcript-level differential usage analysis workflow and downstream consequences classification.

**Table 1.** Cohort design and reciprocal switch results across 13 natural aging tissues.

| **Unit** | **Tissue** | **Cohort (n/group)** | **Aging type** | **n genes tested** | **n tx tested** | **Switches** | **Rate /1k tx** | **Median \|Δπ\|** |
|---|---|---|---|---|---|---|---|---|
| BrainStem | Brain | Female C57BL/6J, 3mo vs 24mo; n□=□5 | Natural | 10,129 | 27,915 | 195 | 6.99 | 0.152 |
| Cerebellum | Brain | Female C57BL/6J, 3mo vs 24mo; n□=□5 | Natural | 9,884 | 27,446 | 156 | 5.68 | 0.140 |
| Cortex | Brain | Female C57BL/6J, 3mo vs 24mo; n□=□5 | Natural | 10,002 | 27,644 | 156 | 5.64 | 0.148 |
| Hippocampus | Brain | Female C57BL/6J, 3mo vs 24mo; n□=□5 | Natural | 10,143 | 28,029 | 121 | 4.32 | 0.147 |
| Midbrain_F | Brain | Female C57BL/6J, 3mo vs 24mo; n□=□5 | Natural | 10,206 | 28,217 | 186 | 6.59 | 0.158 |
| Olfactory | Brain | Female C57BL/6J, 3mo vs 24mo; n□=□5 | Natural | 10,157 | 28,119 | 179 | 6.37 | 0.175 |
| Thalamus | Brain | Female C57BL/6J, 3mo vs 24mo; n□=□5 | Natural | 10,157 | 28,123 | 175 | 6.22 | 0.176 |
| Midbrain_yg_vs_old | Midbrain | Independent cohort; n□=□4 | Natural | 11,327 | 46,373 | 233 | 5.02 | 0.126 |
| Liver_wt2y | Liver | Male, wt8wk ref vs wt2y; n□=□3 | Natural | 6,388 | 15,923 | 134 | 8.42 | 0.234 |
| Liver_wt1.5y | Liver | Male, wt8wk ref vs wt1.5y; n□=□3 | Natural | 6,237 | 15,520 | 86 | 5.54 | 0.238 |
| Adipose_Epi | Adipose | Male epididymal; n□=□3 | Natural | 10,335 | 40,028 | 210 | 5.25 | 0.149 |
| Adipose_Ing | Adipose | Male inguinal; n□=□3 | Natural | 10,564 | 42,459 | 121 | 2.85 | 0.159 |
| Adipose_Mes | Adipose | Male mesenteric; n□=□3 | Natural | 10,887 | 43,393 | 416 | 9.59 | 0.159 |

**Table 2.**
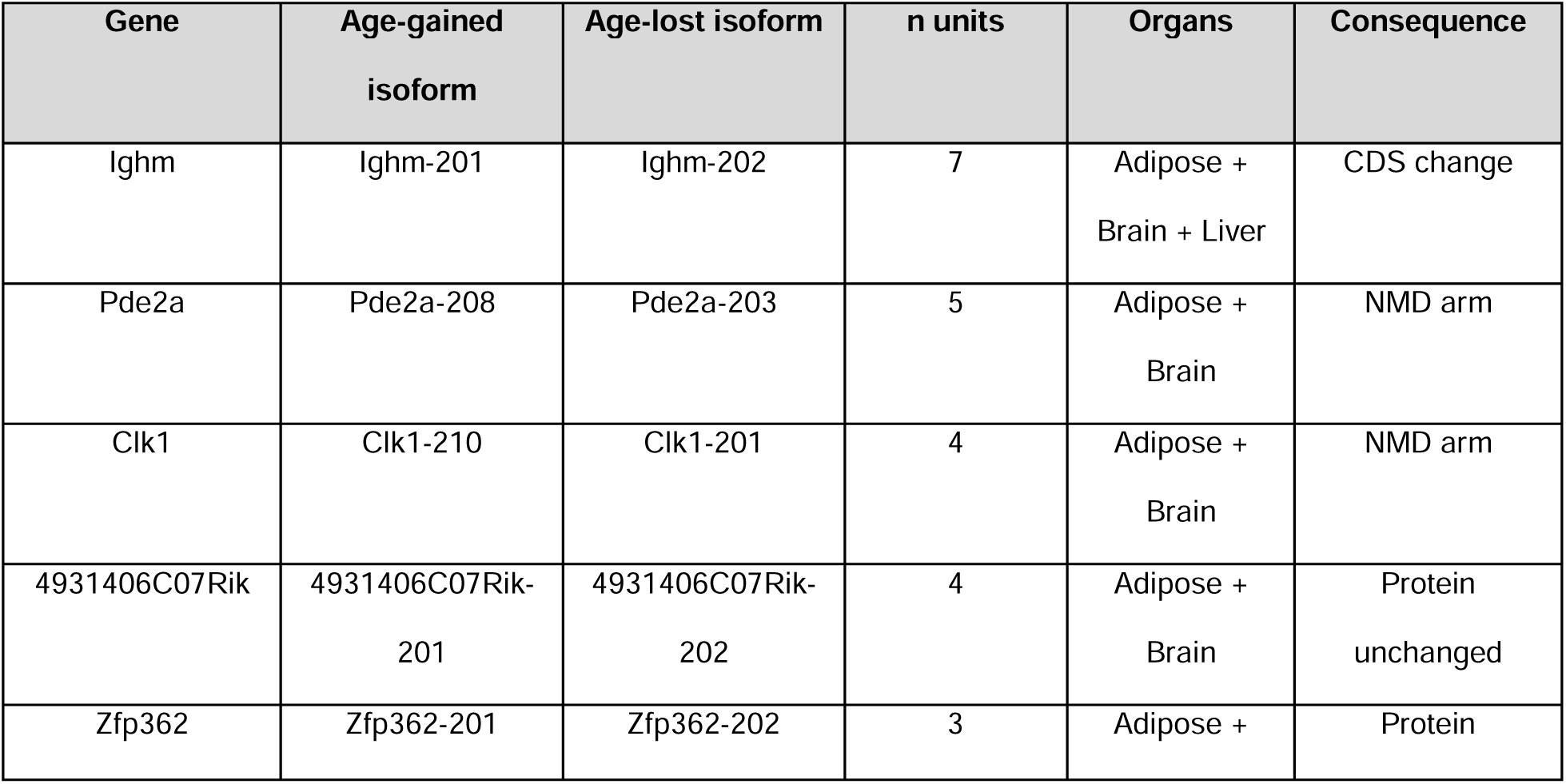
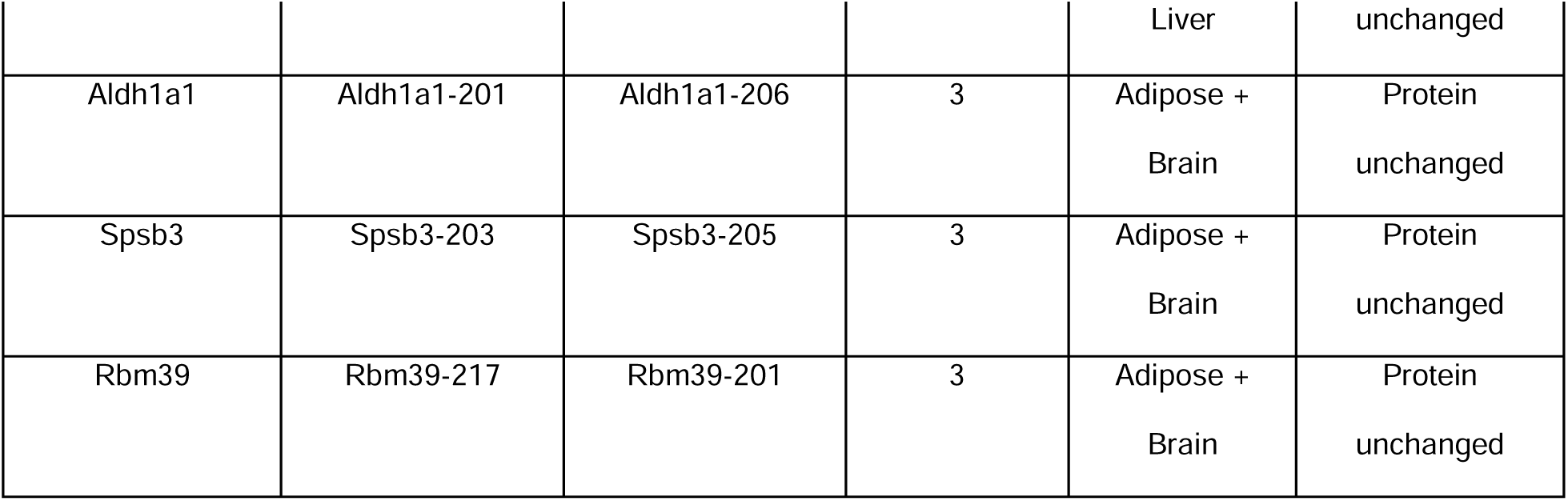
Transcript pairs recurring across two or more organ systems in natural-aging contrasts. Organs = distinct organ systems (brain, liver, adipose) in which the switch is detected. n units = number of individual tissue units. Same pair = whether the identical gained/lost transcript pair is detected in all units. Consequence class from annotation-matched classification.

Within genes showing significant differential transcript usage, we identified transcript pairs in which one isoform increased in relative abundance while another decreased. These reciprocal switches required a change of at least five percentage points in each isoform’s contribution to total gene expression, with at least one isoform reaching transcript-level significance. Across the natural-aging comparisons, we identified 2,368 switches involving 1,779 genes. Of these, 1,168 occurred across the seven brain regions, 233 in the separate midbrain cohort, 220 in liver and 747 in adipose tissue.

To examine whether natural and accelerated aging were associated with similar isoform changes, we separately analyzed Ercc1−/Δ cohorts.

### Age-associated DTU is pervasive across tissues but not uniformly tissue-specific

The number of reciprocal switches per comparison ranged from 86 in the 1.5-year liver contrast to 416 in mesenteric adipose (Table 1). The detected counts reflect the cohort designs, transcripts tested, and significance procedures used in each comparison.

Of the 1,779 distinct switch genes, 1,362 (76.6%) were detected in one comparison, 292 (16.4%) in two, and 125 (7.0%) in three or more. These counts use stable gene identifiers and count each gene once across all 13 comparisons (Supplementary Table 1). We next asked whether recurrence involved the same ordered transcript pair across organs.

We classified predicted coding and untranslated-region consequences separately (Fig. 3a, b)

**Figure 2.**
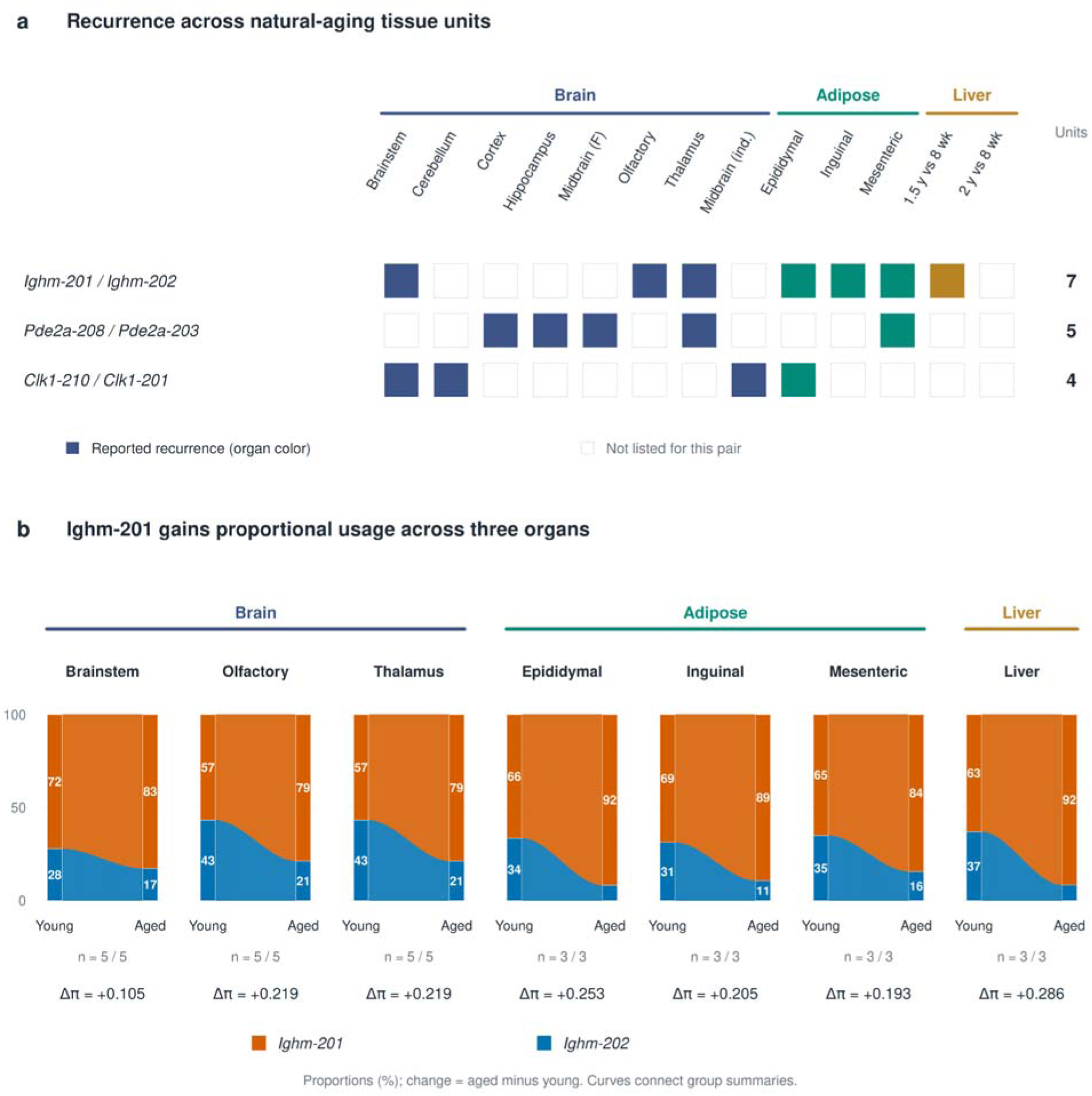
Cross tissue reoccurrence of age associated isoforms switches in Ighm-201 isoform. Fig2a. Distribution of highlighted Ighm, PDe2a and Clk1 transcript pairs across 13 natural aging tissue comparisons. Fig2b. Mean relative isoform proportion of Ighm-201 and Ighm-202 in young and aged groups across seven comparisons in brain, adipose and liver tissues. Ighm-201 usage increases in every comparison (Δπ = 0.105–0.286).

**Figure 3.**
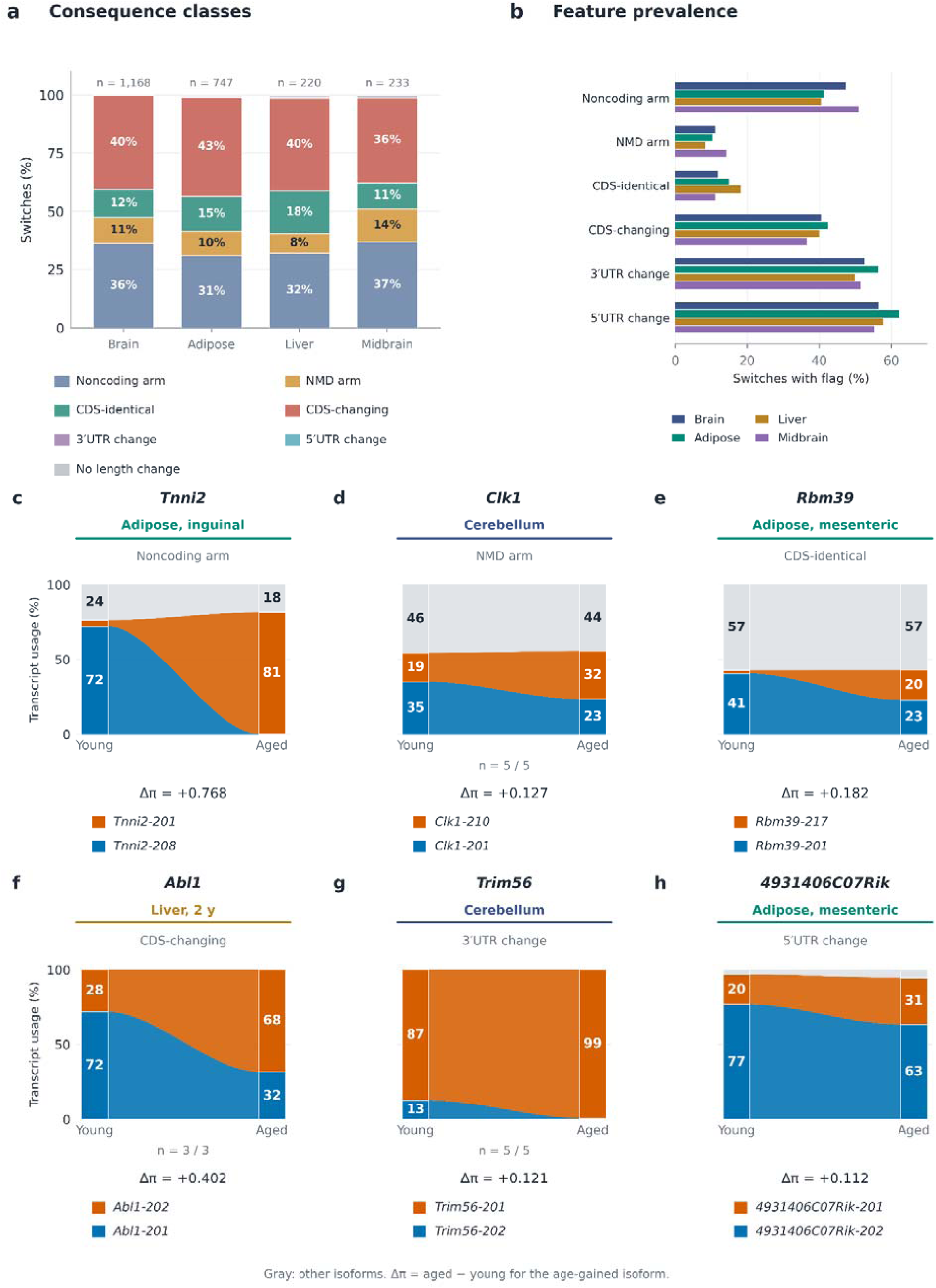
Predicted consequences of age-associated isoform switching. **a.** Proportions of switches assigned to each priority consequence class across brain, adipose, liver, and the independent midbrain cohort. Classes were assigned in the order NMD arm, noncoding arm, CDS-identical, CDS-changing, 3′UTR change, 5′UTR change, and no length change. Numbers above bars indicate total switches. **b.** Prevalence of the six overlapping consequence flags within each tissue group. **c–f.** Exemplar switches representing the noncoding-arm, NMD-arm, CDS-identical, and CDS-changing classes, respectively. **g,h.** Examples with 3′UTR and 5′UTR changes. Ribbons connect young and aged transcript proportions; orange denotes the age-gained isoform, blue the age-lost isoform, and gray the remaining isoforms. Endpoint labels show percentages, and Δπ indicates the aged-minus-young change in usage of the age-gained isoform.

**Figure 4.**
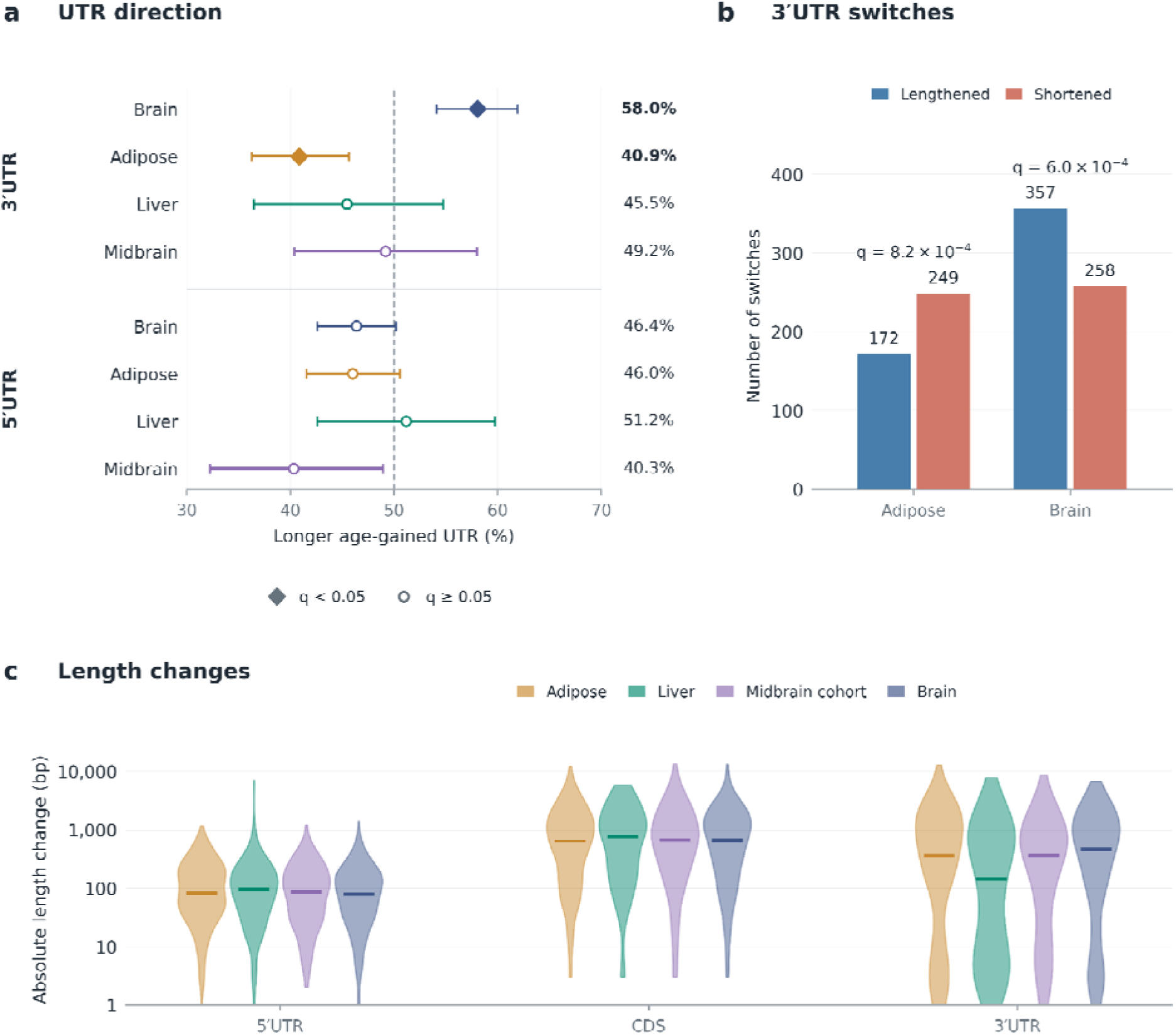
Opposing 3′UTR remodeling in the brain and adipose during aging. **a.** Percentage of switches in which the age-gained isoform has a longer UTR than the age-lost isoform, shown separately for 3′UTRs and 5′UTRs across four tissue groups. Error bars indicate pointwise 95% Wilson confidence intervals; the dashed line marks 50%. Filled diamonds indicate significance after Benjamini–Hochberg correction across eight two-sided binomial tests (q < 0.05). **b.** Numbers of 3′UTR-lengthening and -shortening switches in brain and adipose. **c**.Distributions of absolute 5′UTR, CDS, and 3′UTR length differences across tissue groups, displayed on a logarithmic bp axis. Colored horizontal lines indicate medians; zero-change events are excluded separately for each feature. The brain combines seven female regions; the independent midbrain cohort is shown separately. Counts represent switch events.

### Gene *Ighm* transcript pair recurred across all three organ systems

We defined cross-organ recurrence as detection of the same ordered age-gained and age-lost transcript pair in at least two organ systems. Of the 31 recurring pairs, 23 were shared by brain and adipose only, four by adipose and liver only, and three by brain and liver only. The *Ighm*-201/*Ighm*-202 pair was shared by all three organ systems. It recurred in seven comparisons: all three adipose depots, brainstem, olfactory region, thalamus, and the 1.5-year liver contrast (Fig. 2a, b) . This directional consistency across adipose, brain and liver identifies the Ighm-201/Ighm-202 switch as the most robust age-associated isoform event in the dataset (Fig. 2B). A recent expression-level survey identifies Ighm among the most reproducibly age-upregulated genes across murine tissues^6^; the present data resolve this to isoform resolution, showing the secreted-form Ighm-201 transcript gaining proportional usage at the expense of the membrane-anchored Ighm-202 variant across all three organ systems.

The third most-recurrent same-pair switch, Clk1-210/Clk1-201 (consequence: NMD arm), appeared in four tissue units spanning adipose (Adipose_Epi) and brain (Brain Stem, Cerebellum, and the Midbrain tissue; Δπ range 0.110–0.195). CLK1 is a dual-specificity serine-threonine kinase that phosphorylates the RS (arginine–serine) domains of SR proteins, which regulate exon inclusion and splice-site selection.^7^ The increased relative usage of NMD-annotated *Clk1*-210, accompanied by decreased usage of protein-coding *Clk1*-201, suggests a shift in the balance between potentially productive and NMD-targeted transcripts. Whether this shift affects CLK1 protein abundance, kinase activity, or downstream splicing remains to be established.

The *Pde2a*-208/*Pde2a*-203 switch recurred in five tissue units: four regions of the female brain cohort (cortex, olfactory region, midbrain, and thalamus) and mesenteric adipose tissue. In each unit, NMD-annotated *Pde2a*-208 gained relative usage while protein-coding *Pde2a*-203 lost usage. PDE2A is a dual-substrate phosphodiesterase that hydrolyzes both cAMP and cGMP. This shared directional pattern identifies a recurrent age-associated shift in *Pde2a* transcript usage across brain and adipose tissue. Whether it affects PDE2A protein abundance or cyclic nucleotide signaling remains to be determined.

### Most age-associated isoform switches do not alter protein sequence

We classified each switch pair using six non-exclusive binary flags on annotation-matched transcript models: NMD arm, noncoding arm, CDS-identical, CDS-changing, 3′UTR change, and 5′UTR change. A priority consequence class was assigned in that order. Across the four organ groups (Fig. 3a, b), 31.2–36.5% of switches involve a noncoding arm, an additional 8-14% involve an NMD-annotated arm, and 11-18% are protein-identical when tested by exact CDS exon-coordinate signature. Together, these regulatory and protein-null events account for most of all switches in every tissue. CDS-changing switches constitute 36.5 -42.6% of all events, but only a minority of those meet stricter criteria for structural ensemble modeling.

The protein-identical class (11–18%) is invisible to any rule based on CDS length or domain count, and its prevalence means that a substantial fraction of CDS-involving switches have no protein consequence. Rbm39, a splicing-regulatory factor recurring in three tissue units across Brain and Adipose (Adipose_Mes, Olfactory, Thalamus), belongs to this class: its two isoforms diverge in 5′UTR architecture but encode the same protein a splicing regulator whose own regulation is UTR-mediated. This observation suggests a post-transcriptional auto-regulatory feedback mechanism that would be entirely invisible to exon-level or domain-count approaches.

#### Brain and adipose show opposing 3′UTR remodeling signatures

Among switches that change the 3′UTR, we tested for directional bias in the age-gained isoform across eight tissues and UTR combinations, correcting with Benjamini–Hochberg across all eight tests, using annotation-matched transcript lengths. Two tissues reach significance and point in opposite directions: brain 3′UTR switches preferentially lengthen the age-gained isoform (357 of 615 switches lengthened; 58.0%; q = 6.0×10⁻ ), while adipose switches preferentially shorten it (249 of 421 shortened; 59.1%; q = 8.2×10⁻ ). Liver (45.5% lengthened, n = 110, q = 0.52) and the midbrain cohort (49.2%, n = 120, q = 0.93) are not significant. No 5′UTR directional bias is significant in any tissue (Fig. 5a).

**Figure 5.**
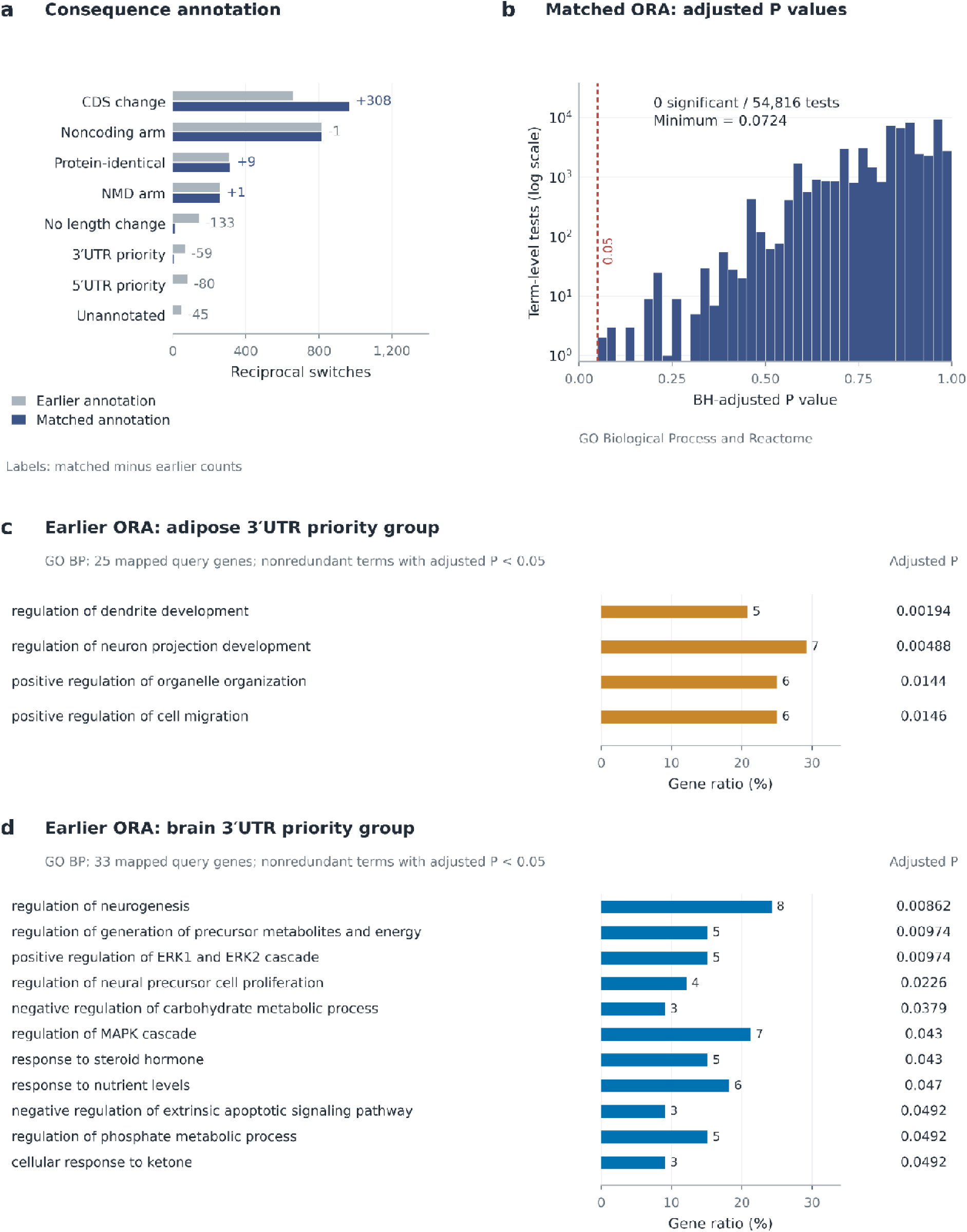
Pathway analysis of age-associated isoform switch genes. **a.** Numbers of GO Biological Process, KEGG, and Reactome terms tested for each organ and consequence group. **b.** Highest-ranked enrichment results for the complete adipose switch-gene set, using genes tested for differential transcript usage as the background. Values show −log₁₀-transformed Benjamini-Hochberg-adjusted enrichment p-values. Highlighted points denote splicing-related terms identified by keyword filtering.

The magnitude of 3′UTR changes in the significant tissues is substantial: individual events reach several kilobases of added or subtracted sequence, with the distribution centering near 1,000 bp (Fig. 5b). The largest brain 3′UTR extension is Trim56 in Cerebellum (+6,812 bp) and Fam204a in Midbrain (+5,804 bp); the largest adipose shortening includes Prlr in Adipose Mes (−7,742 bp).

Among switches with a 3′ UTR length change, the age-gained transcript had a longer 3′ UTR in 58.0% of switches pooled across the seven female brain regions and in 49.2% of switches in the independent midbrain cohort. Differences in cohort design and reference annotation limits direct comparison. Comparing the two midbrain datasets after reprocessing them with a common reference would provide a closer comparison, while estimates for each female brain region would show how consistently the lengthening bias was observed across regions.

### Age-associated switch genes show no pathway-level enrichment

Annotation-matched pathway analysis detected no significant over-representation of GO Biological Process terms or Reactome pathways among switch genes in brain or adipose tissue. We examined both the complete switch-gene set in each organ and six overlapping groups defined by the predicted consequences of their isoform switches. Each set was compared with the genes tested for DTU in the corresponding organ. None of the 54,816 term-level tests met the threshold of BH q < 0.05, with correction applied separately for each organ, consequence group, and database. The lowest adjusted value was q = 0.072 for the Reactome terms “Cell junction organization” and “Cell-Cell communication” in the complete adipose switch-gene set (Fig. 6b).

**Fig. 6.**
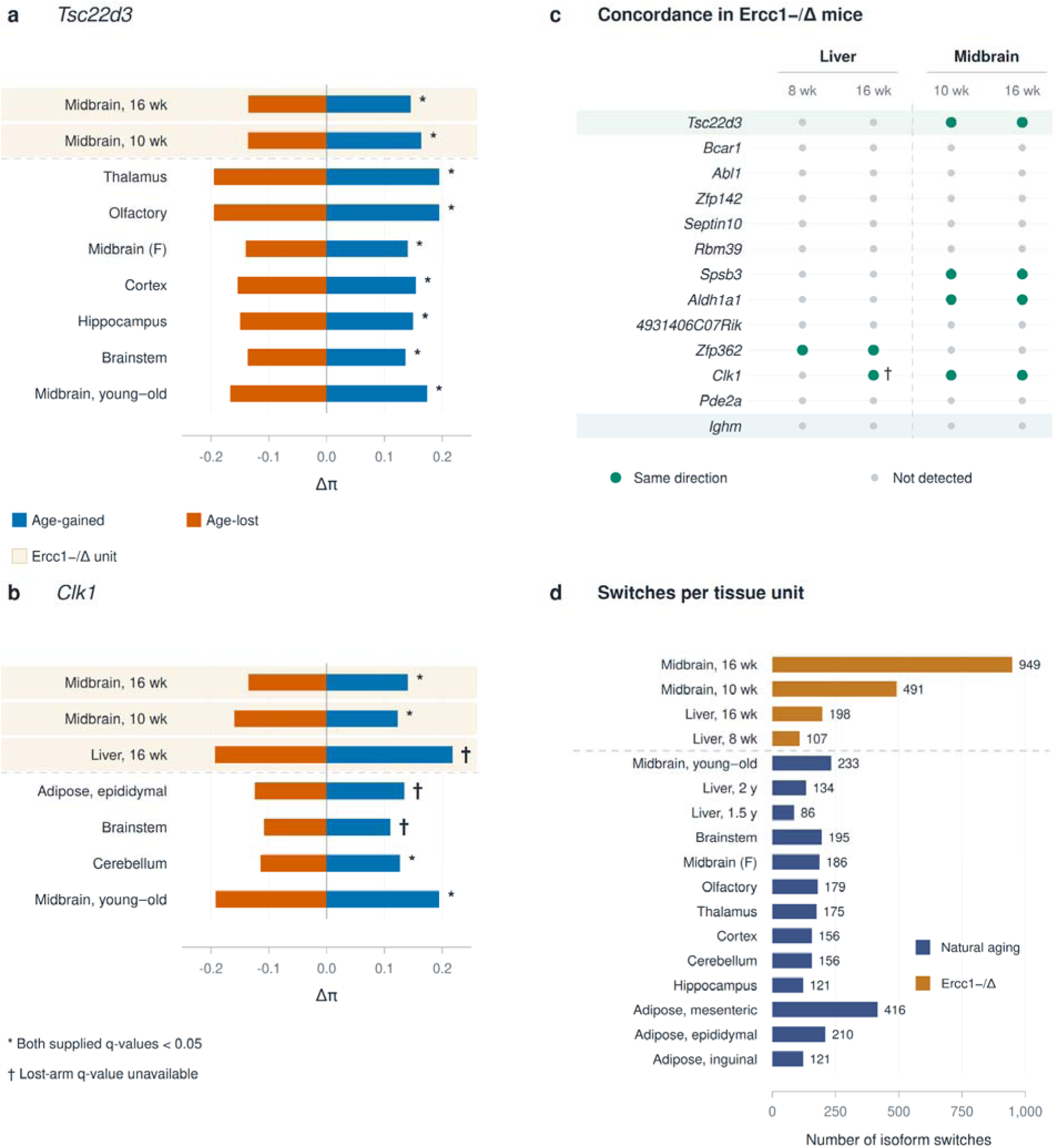
Ercc1−/Δ mice partially recapitulate the natural aging isoform-switch landscape. a,b. Δπ (proportional isoform usage shift, young versus aged) for the age-gained (blue) and age-lost (red) isoform arms of *Tsc22d3* (a) and *Clk1* (b) across natural aging (white) and Ercc1−/Δ (shaded) tissue units. Asterisks indicate units in which both isoform arms reach significance (q < 0.05). **c.** Concordance of the 13 most recurrent natural aging isoform switches across four Ercc1−/Δ tissue units. Green circles, same direction as natural aging; red triangles, reversed direction; grey circles, not detected. **d**. Number of significant isoform switches per tissue unit in natural aging (blue) and Ercc1−/Δ mice (orange). Colored dots indicate organs of origin. Dashed lines separate natural aging from Ercc1−/Δ units.

**Figure 7.**
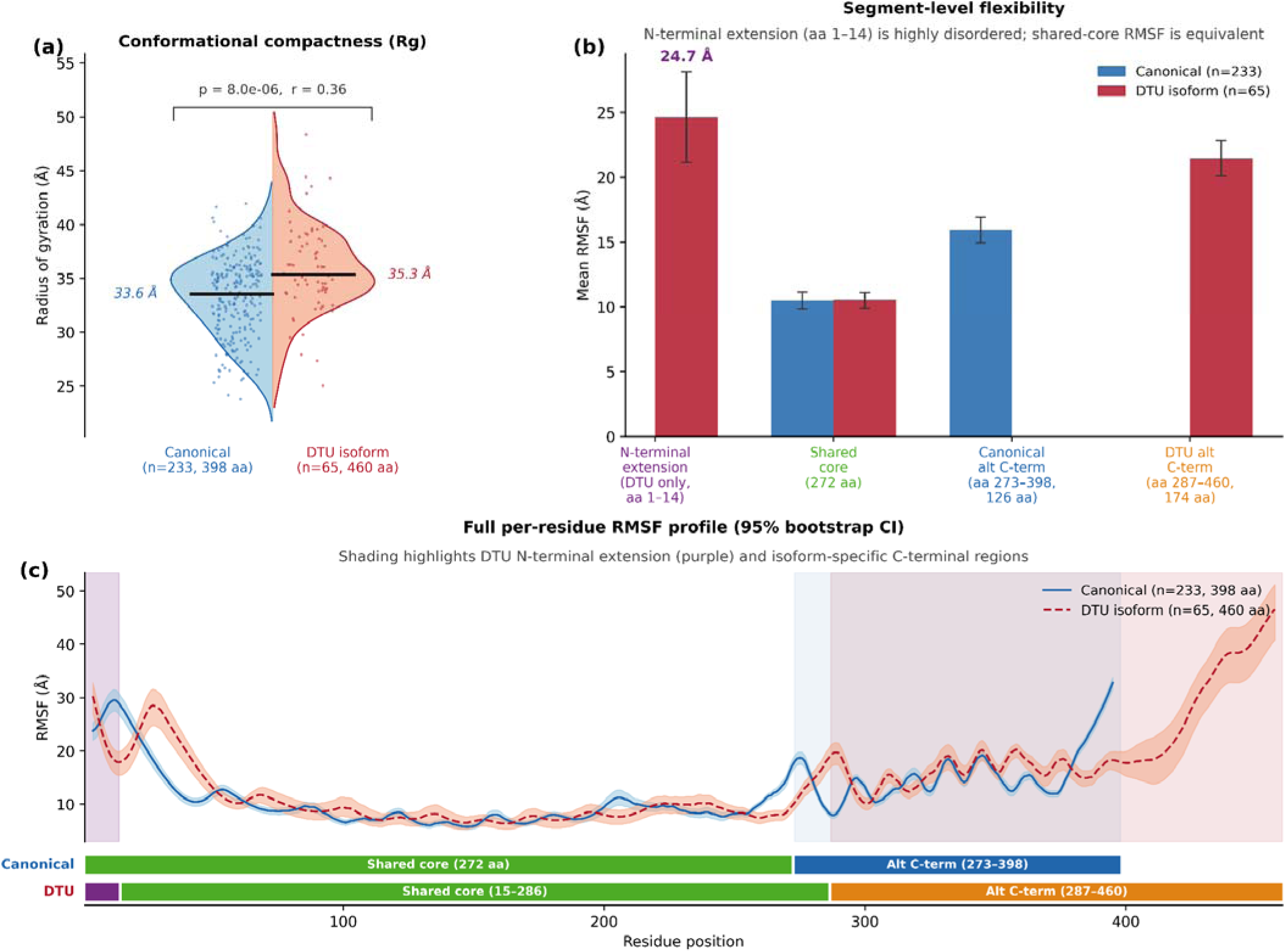
Conformational properties of *Cyth1* isoforms modeled with BioEMU. **a.** Radius-of-gyration distributions for the canonical and age-associated isoform ensembles. Points represent individual conformations; distributions were compared using the Mann– Whitney U test. **b.** Mean root mean square fluctuation (RMSF) of the N-terminal extension, shared core, and isoform-specific C-terminal regions. Error bars indicate 95% confidence intervals. **c.** Per-residue RMSF profiles following iterative Kabsch superposition, with shaded bands indicating 95% bootstrap confidence intervals. Background shading and sequence tracks identify the shared core and isoform-specific regions. Blue denotes the canonical isoform and red the age-associated isoform.

We also examined splicing-related terms within the pathway enrichment results. Keyword filtering identified 25 distinct term descriptions, represented by 255 tests across organs and consequence groups; none was significant after correction (minimum q = 0.46). Combining brain and adipose enrichment p-values using Fisher’s method produced the smallest splicing-related p-value for “regulation of RNA splicing” among genes with switches involving an NMD-annotated transcript (combined p = 0.0016). This result did not remain significant after BH correction across 21,029 matched term–consequence tests (combined q = 0.31).

### Ercc1−/Δ mice aging partially recapitulates the natural aging switch landscape

The Ercc1−/Δ model provides an opportunity to test whether accelerated aging engages the same isoform-switching landscape as chronological aging. Among the recurrent multi-unit switches, Tsc22d3 and Clk1 both agree in direction between natural and progeroid units. Tsc22d3 switches in the same direction in both Midbrain_ercc10wk and Midbrain_ercc16wk progeroid units as in the natural aging brain cohort, suggesting that glucocorticoid receptor signaling changes at the isoform level are a conserved feature of both aging modalities. Clk1, which recurs in natural aging across Adipose Epi, Brain Stem, Cerebellum, and Midbrain, also agrees in direction in Liver_ercc16wk, Midbrain_ercc10wk, and Midbrain_ercc16wk progeroid units. Ighm, the pan-organ switch and the most recurrent event in natural aging, is absent from all Ercc1−/Δ models.

### A sequence-level eligibility cascade limits structural interpretation to tissue-restricted events

We screened 35 reciprocal isoform switches detected in at least three brain regions for conformational ensemble modeling. Seventeen pairs remained after excluding switches in which either transcript was noncoding or predicted to undergo nonsense-mediated decay. Requiring a protein sequence difference of at least 30 amino acids reduced the set to 13 pairs. Eight pairs remained after requiring that the longer isoform be no more than twice the length of the shorter one. To limit computational demands, we restricted BioEMU sampling to pairs in which both isoforms were 600 amino acids or shorter, leaving five eligible pairs. Song et al. used a similar length cutoff for large-scale AlphaFold2 modeling of human splice isoforms. The coding and nonsense-mediated decay criteria accounted for the largest reduction, excluding 18 of the 35 switches.

To our knowledge, this is the first application of equilibrium ensemble emulation to age-associated isoform switches. We applied BioEMU^10^ to generate 300 conformational samples for the eligible Cyth1 canonical isoform (398 aa, SEC7+PH domains) and the DTU isoform (460 aa, alternative first exon adding a 14-aa N-terminal extension (the two isoforms differ by 62 aa in total: 14 aa gained at the N-terminus plus a distinct 174-aa C-terminus vs the canonical 126-aa C-terminus)). The full-length DTU isoform (460 aa) has significantly larger radius of gyration than the canonical form (398 aa) (canonical median 33.6 Å vs. DTU median 35.4 Å; Mann– Whitney U p = 8.0×10⁻ , rank-biserial r = 0.36), consistent with the 14-aa disordered N-terminal extension and larger alternative C-terminus expanding the conformational envelope. The shared 272-residue core (canonical aa 1–272; DTU aa 15–286) shows strongly correlated per-residue RMSF (Pearson r = 0.982, p < 10⁻¹ , n = 272 residues) (Fig. 8b). Divergence is confined to the disordered N-terminal extension (DTU aa 1–14; mean RMSF 24.7 Å; 95% CI 21.5–28.2 Å) and the isoform-specific C-terminal regions (DTU: 21.4 Å, 95% CI 20.1–22.8 Å; canonical: 15.9 Å, 95% CI 15.0–16.9 Å), relative to the shared core (10.5 Å). The alternative N-terminal exon of the DTU isoform is intrinsically disordered and does not change the SEC7 and PH domain fold, suggesting that the aging-gained Cyth1 isoform preserves the catalytic and membrane-interaction functions of the canonical protein while acquiring a flexible N-terminal appendage whose regulatory role, if any, remains to be determined.

### Validation gap and experimental limitations

No isoform-level validation by RT-PCR was performed. Isoform-specific RT-qPCR validation targeting Tsc22d3 (7-unit brain switch), Ighm (pan-organ), a protein-identical switch (Nup37 or Fgr), and a 3′UTR-lengthening event (Trim56 or Fam204a) will form the basis of future experimental work. Until validated, the switches reported here should be treated as statistically supported computational predictions requiring orthogonal confirmation.

## Discussion

Age-associated isoform remodeling across brain, liver and adipose tissue involves changes in both transcript regulation and protein coding potential. In this study, we identified 2,368 switches across 1,779 genes across three tissues. Most switches involved at least one noncoding transcript, an isoform annotated as a target of nonsense-mediated decay (NMD), or transcripts with different structures that encode identical proteins. These findings identify a substantial component of age-associated remodeling whose predicted consequences include transcript fate or the expression between coding and noncoding isoforms. They complement the observation by Sieber et al.^2^ that protein domain counts are largely preserved during aging, while showing why domain-level comparisons alone cannot capture the full range of transcript changes. However, these annotations identify potential consequences; they do not establish changes in RNA stability, translation, or protein abundance.

The observed biological signal in this dataset is the tissue opposing directionality of 3′UTR remodeling in brain and adipose. The brain systematically gains longer 3′UTRs with age (58.0%; q = 6.0×10⁻ ), while adipose systematically shortens them (59.1%; q = 8.2×10⁻ ). This pattern is consistent with established associations between alternative polyadenylation (APA) and cellular state: shorter 3′UTRs are frequently associated with proliferation and immune activation, whereas longer 3′UTRs accompany differentiation in several experimental systems^11–13^. These associations provide a framework for interpreting our findings, but they do not establish that the aging tissues follow the same underlying mechanism.

The opposing 3′UTR patterns in brain and adipose suggest that age-associated isoform remodeling may have different post-transcriptional consequences across tissues. Extended 3′UTRs can contain additional microRNA and RNA-binding-protein recognition sites, creating opportunities for regulation beyond the protein-coding sequence^13^. Experimental studies have demonstrated that alternative 3′UTRs can influence RNA localization and stability, although these effects depend on the gene, UTR sequence, and cellular context^14,15^. The preference for longer isoforms in the brain and shorter isoforms in adipose could therefore alter the regulatory interactions available to these transcripts. Our analysis does not establish which interactions are affected or whether the observed patterns arise within cells. Comparing the gained and lost UTR segments for enrichment of specific regulatory motifs, using backgrounds matched for sequence length and composition, would help prioritize mechanisms for experimental testing.

The *Ighm* transcript pair was the only switch shared across all three organ systems, with the same direction of change observed in seven tissue comparisons. This recurrence extends previous reports of age-associated *Ighm* upregulation across murine tissues^6^ by identifying a shared shift in relative transcript usage toward *Ighm*-201, annotated as encoding the secreted IgM heavy chain form. The distinction is biologically relevant because secreted and membrane-bound immunoglobulin heavy chains are generated through alternative RNA processing linked to B-cell differentiation^16^.

Age-associated changes in human B-cell function and subset composition^17,18^, together with plasma-cell accumulation in aging mice^19^, provide a plausible context for the shared *Ighm* switch. The recurrence of *Jchain* switches involving a noncoding transcript in five brain comparisons is compatible with an immune-associated origin, although it does not establish increased JCHAIN protein production or infiltration. Increased B-cell abundance alone would not necessarily alter *Ighm* transcript proportions unless the contributing populations differed in isoform usage. The shared switch could therefore reflect changes in the balance of B-cell and antibody-secreting populations, altered RNA processing within these cells, or both.

Experimental validation through qRT-PCR is needed to confirm these switches and determine the reflected changes in altered RNA processing of all the tissues.

Reciprocal isoform switches showed no significant enrichment for biological processes or pathways in brain or adipose, including when analyzed separately by coding potential, coding sequence or UTR length change. Targeted analysis of splicing-related functions and the combined analysis across organs likewise showed no significant enrichment after multiple-testing correction. These findings provide no evidence that the identified switches preferentially affect genes involved in the splicing machinery. Individual switches may nevertheless have biological consequences that gene-level pathway analysis does not capture, particularly when transcript differences affect regulatory sequences or protein structure. The absence of significant enrichment therefore does not exclude functional effects, but limits conclusions about coordinated involvement in particular pathways.

The Ercc1−/Δ comparison provides partial evidence of overlap with the natural aging program: Tsc22d3 and Clk1 agree in direction in the Ercc1−/Δ mice model, while Ighm is absent. The incomplete concordance is consistent with prior gene-expression analyses in the same model and argues that Ercc1−/Δ mice captures a component of the aging transcriptomic landscape while also engaging mechanisms not shared with chronological aging. Whether Clk1’s recurrence across both aging modalities reflects a conserved post-transcriptional mechanism or a shared upstream stress response remains to be determined.

Bulk-tissue RNA sequencing captures isoform changes across mixed cell populations, limiting our ability to distinguish regulation within individual cell types from changes in cellular composition. Short-read quantification also introduces uncertainty when transcripts share extensive sequence, particularly for closely related isoforms. Furthermore, transcript annotations and structural ensemble predictions identify potential functional consequences but do not establish their effects in aging tissues. Experimental studies will be needed to determine whether selected switches alter RNA stability, translation, protein abundance, or cellular function. These caveats define the experimental work required to confirm the computational findings reported here to the level of validated biological functions.

Together, these findings establish that age-associated isoform switching is a widespread feature of the aging transcriptome but operates predominantly through non-protein-altering mechanisms. The tissue-opposing 3′UTR remodeling in brain and adipose defines a candidate layer of post-transcriptional regulatory divergence between two organs that age in markedly different cellular contexts. The structural eligibility cascade described here provides a principled template for prioritizing ensemble modeling resources for any DTU discovery dataset, and the Cyth1 BioEMU example demonstrates the feasibility of conformational characterization of aging-associated isoform pairs.

## Methods

### Animals and tissue collection

Brain tissue comprised seven regions (brainstem, cerebellum, cortex, hippocampus, midbrain, olfactory bulb, and thalamus) from female C57BL/6J mice aged 3 or 24 months (n = 5 per age group). An independent midbrain cohort included young and aged mice. The primary natural-aging liver comparisons used male mice aged 1.5 and 2 years and an 8-week-old reference group (n = 3 per group). The adipose analysis included epididymal, inguinal, and mesenteric depots. The Ercc1−/Δ mice comparisons included midbrain at 10 and 16 weeks and liver at 8 and 16 weeks

### RNA-seq processing and transcript quantification

RNA-seq processing used STAR and Salmon with cohort-specific reference genomes and annotations.^20,21^ The seven female brain regions used GRCm39 with Ensembl release 110. The matched analysis used GRCm38-associated annotations for the other cohorts. Salmon estimates were imported with tximeta.^22^

### Differential transcript usage analysis

We tested for differential transcript usage using DRIMSeq, which models transcript proportions with a Dirichlet–multinomial distribution.^23^ A design matrix specifying age group as the only factor was used for all comparisons. DRIMSeq gene-level and transcript-level p-values were passed to stageR for stage-wise false discovery rate control at α = 0.05 across all comparisons.^24^ A transcript was considered significant if the gene cleared the screening stage (stageR q_gene < 0.05) and the transcript cleared the confirmation stage (stageR q_tx < 0.05). DRIMSeq and stageR were run in R version 4.3.x (Bioconductor 3.18).

### Definition of reciprocal switches

A reciprocal switch was defined as a gene satisfying all of the following criteria: (1) gene-level significance as defined above; (2) at least one transcript arm reaching transcript-level significance under the same framework; (3) at least one transcript with Δπ > 0 (gaining arm) and at least one with Δπ < 0 (losing arm), where Δπ = π aged − π young (the DRIMSeq maximum-likelihood estimate of isoform proportion within the gene in the aged and young groups, respectively); and (4) both the gaining and losing arm satisfying |Δπ| ≥0.05. Among genes meeting criteria 1–4, the gaining arm is the transcript with the largest positive Δπ and the losing arm is the transcript with the most negative Δπ. Switches were identified per unit independently; no cross-unit multiplicity correction is applied. Switch rate was calculated as the number of reciprocal switches divided by the number of tested transcripts, multiplied by 1,000.

### Transcript consequence classification

We classified switches using annotation-matched transcript models and six overlapping flags for a noncoding arm, an NMD-annotated arm, matching CDS exon coordinates, different CDS exon coordinates in coding and non-NMD pairs, and differences in 3′UTR or 5′UTR length. We assigned one priority class per switch in this order: NMD arm, noncoding arm, CDS-identical, CDS-changing, 3′UTR change, 5′UTR change, and no length change. Identical CDS signatures required the same chromosome, strand, and ordered coding-exon coordinates.

### UTR directional analysis

For each tissue group and UTR type, we compared the annotation-matched UTR length of the gaining and losing transcripts. We excluded switches with no length difference for that UTR type and used two-sided binomial tests against a lengthening fraction of 0.5. We applied Benjamini– Hochberg correction across all eight tests. Brain pooled the seven female regions, adipose pooled the three depots, and liver pooled the 1.5 and 2-year contrasts. The independent midbrain cohort was analyzed separately. Counts represent switch events.

### Pathway over-representation analysis

Switch genes per organ were tested for pathway over-representation against the genes tested for DTU in that organ (background universe). Gene symbols were mapped to Entrez IDs using the org.Mm.eg.db annotation package (96.3% mapping rate; 13,853 of 14,388 unique tested genes). Over-representation analysis used clusterProfiler::enricher^25^ applied to GO Biological Process terms^26^, KEGG pathways^27^, and Reactome pathways^28^. Tests were run separately for each of six consequence strata (non_coding_arm, nmd_arm, cds_changed, cds_identical, utr3_changed, utr5_changed) and for the combined ALL stratum in brain and adipose. BH correction was applied within each stratum × organ combination. A total of 54,816 terms were tested across all combinations. Spliceosome replication was assessed by extracting 204 GO, KEGG, and Reactome terms containing the keywords ‘spliceosom’, ‘splicing’, or ‘RNA processing’ and applying the same over-representation test; no term reached significance (best q = 0.46).

### BioEMU structural ensemble analysis

We screened reciprocal isoform switches detected in at least three brain regions for structural ensemble modeling. Both transcripts had to be protein coding and neither could be predicted to undergo nonsense-mediated decay. We required the encoded proteins to differ by at least 30 amino acids, the longer isoform to be no more than twice the length of the shorter isoform, and both isoforms to be 600 amino acids or shorter. We retrieved protein translations through the Ensembl REST API using the annotation release matched to each transcript analysis.

For the original *Cyth1* comparison, we generated 300 conformations per input sequence using BioEMU¹ with default parameters. After filtering, 233 conformations remained for the 398-amino-acid canonical isoform and 65 for the 460-amino-acid alternative isoform. We calculated the radius of gyration for each conformation as the root-mean-square distance of Cα atoms from their centroid. To calculate per-residue root-mean-square fluctuation (RMSF), we aligned conformations to a common mean structure using three iterations of Kabsch superposition, with residues 1–377 of the canonical isoform and 15–391 of the alternative isoform as alignment anchors. We compared radius-of-gyration distributions using a two-sided Mann–Whitney U test and estimated 95% confidence intervals for RMSF using 1,000 bootstrap resamples with replacement.

### Statistics

All statistical analyses were performed in R (version 4.3.x). DTU testing used DRIMSeq and stageR as described above. Binomial tests used the base R binom.test function. Multiple testing corrections used the Benjamini–Hochberg procedure throughout. Group comparisons used the Mann–Whitney U test (wilcox.test, two-sided). Cohen’s d was computed as the standardized mean difference using the pooled standard deviation. Bootstrap confidence intervals used the boot package. Pearson correlation coefficients and their p-values were computed with cor.test. No statistical power calculation was performed; sample sizes are determined by the original experimental designs.

## Funding Acknowledgments

The work reported here received financial support from NIH grants U54 AG079754 and U54 AG076041. The UMN Masonic Cancer Center is supported by the U.S. National Institute of Health and National Cancer Institute [Cancer Center Support Grant CA-77598].

## Author Contributions

J.W., L.J.N., and P.D.R. conceived and designed the study. N.P., J.W., and S.A. carried out the data collection and the data analysis. N.P., L.L., A.H., M.D., E.K., C.A., P.D.R., L.J.N., J.W. contributed to data interpretation. N.P., and J.W. wrote the manuscript, and J.W. reviewed and edited the manuscript with feedback from all the authors. All authors reviewed and approved the final manuscript.

