## Supplemental Table 1 for "A multi-tissue atlas of age-associated differential transcript usage in mouse reveals predominantly regulatory isoform switching across brain, liver, and adipose"

Supplementary Table 1 | Counts and recurrence of age-associated isoform switches

The table summarizes reciprocal isoform switches across 13 natural-aging comparisons. Each switch comprises an age-gained transcript and an age-lost transcript of the same gene within a comparison.

a Switch counts by tissue group

| **Tissue group** | **Comparisons** | **Switches** | **Distinct genes** |
| --- | --- | --- | --- |
| Female brain regions | 7 | 1,168 | 843 |
| Independent midbrain | 1 | 233 | 233 |
| Liver | 2 | 220 | 210 |
| Adipose | 3 | 747 | 694 |
| **Overall** | **13** | **2,368** | **1,779** |

b Gene recurrence across comparisons

| **Comparisons with a switch** | **Distinct genes** | **Share of all switch genes** |
| --- | --- | --- |
| 1 | 1,362 | 76.6% |
| 2 | 292 | 16.4% |
| 3 | 92 | 5.2% |
| 4 | 23 | 1.3% |
| 5 | 8 | 0.4% |
| 6 | 0 | 0.0% |
| 7 | 2 | 0.1% |

Genes were counted by stable gene identifier. Panel a counts distinct genes within each tissue group, so a gene can appear in more than one group. The overall row counts each gene once across all comparisons. Panel b counts each gene once according to the number of comparisons in which it switched, regardless of whether the same transcript pair recurred. Percentages use all 1,779 distinct switch genes as the denominator.
